# ATM inhibition drives metabolic adaptation via induction of macropinocytosis

**DOI:** 10.1101/2020.04.06.027565

**Authors:** Chi-Wei Chen, Raquel Buj, Erika S. Dahl, Kelly E. Leon, Erika L. Varner, Eliana von Krusenstiern, Nathaniel W. Snyder, Katherine M. Aird

## Abstract

Macropinocytosis is a nonspecific endocytic process that enhances cancer cell survival under nutrient-poor conditions. Ataxia-Telangiectasia mutated (ATM) is a tumor suppressor that plays a role in cellular metabolic reprogramming. We report that suppression of ATM increases macropinocytosis in an AMPK-dependent manner to promote cancer cell survival in nutrient-poor conditions. Combined inhibition of ATM and macropinocytosis suppressed proliferation and induced cell death both *in vitro* and *in vivo*. Metabolite analysis of the ascites and interstitial fluid from tumors indicated decreased branched chain amino acids (BCAAs) in the microenvironment of ATM-inhibited tumors. Supplementation of ATM inhibitor-treated cells with BCAAs abrogated AMPK phosphorylation and macropinocytosis and rescued the cell death that occurs due to combined inhibition of ATM and macropinocytosis. These data reveal a novel molecular basis of ATM-mediated tumor suppression whereby loss of ATM promotes pro-tumorigenic uptake of nutrients to promote cancer cell survival and reveal a metabolic vulnerability of ATM-inhibited cells.

## Introduction

Macropinocytosis is an endocytic process whereby cells take up fluid, macromolecules, metabolites, and other cargo from the surrounding microenvironment (King and Kay, 2019; Palm, 2019; Zhang and Commisso, 2019). Many studies have evaluated the critical role of macropinocytosis as a nutrient scavenging mechanism in cancers under nutrient-deprived conditions (Commisso et al., 2013; Davidson et al., 2017; Hodakoski et al., 2019; Kamphorst et al., 2015; Kim et al., 2018; Lee et al.; Redelman-Sidi et al., 2018; Tejeda-Munoz et al.; Yao et al., 2019). To date, these studies have focused on cancers with high PI3K activity, such as those with mutant RAS or PTEN loss, which act upstream of the Rac1-Pak1 actin remodeling pathway to promote macropinosome formation (Zhang and Commisso, 2019). As macropinocytosis is a process that is driven by multiple nutrient sensing pathways, including AMPK and mTOR (Kim et al., 2018; Palm et al., 2015), it is likely that other tumor-associated pathways also induce macropinocytosis as a means to provide nutrients.

Ataxia-Telangiectasia mutated (ATM) is a tumor suppressor, and mutation or loss of ATM expression promotes genomic instability and predisposes cells to tumorigenesis (McKinnon, 2004, 2012). While ATM is known to be important for the response to DNA double strand breaks (Shiloh, 2003), it is also known to play a role in cellular metabolism (Dahl and Aird, 2017; Guleria and Chandna, 2016). We and others have shown that inhibition of ATM increases uptake of glucose and glutamine to provide nutrients for cell growth and proliferation (Aird et al., 2015; Dahl and Aird, 2017). Our previous results suggest that inhibition of ATM reprograms metabolism through suppression of p53 signaling and enhanced c-MYC protein stability (Aird et al., 2015). Indeed, these and other models where ATM suppression or mutation has been shown to alter cellular metabolism are in cells with wildtype p53 and normal c-MYC expression. Whether ATM similarly alters nutrient uptake and cellular metabolism in the context of cancers with mutated p53 and high c-MYC expression is unclear. Moreover, ATM inhibitors are currently undergoing clinical trials (Jin and Oh, 2019); thus, understanding how inhibition of ATM drives metabolic reprogramming may be important towards identifying potential resistance mechanisms or other targets that could be used in combination with these inhibitors in ATM-wildtype tumors.

We found that suppression of ATM increases nutrient uptake in p53 mutant and c-MYC amplified cancer cells independently of transporters. This non-specific nutrient uptake under nutrient poor conditions occurred via AMPK-dependent macropinocytosis. Underscoring the importance of macropinocytosis in ATM-inhibited cells, suppression of macropinocytosis using 5-(N-ethyl-N-isopropyl) amiloride (EIPA) (Commisso et al., 2014; Ivanov, 2008) significantly inhibited proliferation of cancer cells both *in vitro* and *in vivo*. Analysis of tumor metabolites suggested increased uptake of branched chain amino acids (BCAAs) in ATM inhibited tumors, and supplementing cells *in vitro* with exogenous BCAAs decreased macropinocytosis and AMPK phosphorylation and suppressed the synergy between ATM and macropinocytosis inhibition. Together, these data demonstrate that inhibition of ATM reprograms cellular metabolism through AMPK-mediated induction of macropinocytosis and reveal that macropinocytosis is a vulnerability of ATM inhibited tumors.

## Results and Discussion

### Inhibition of ATM kinase activity increases glucose and glutamine consumption in a transporter-independent manner

ATM loss or inhibition alters whole body and cellular metabolism (Aird et al., 2015; Cosentino et al., 2011; Dahl and Aird, 2017; Guleria and Chandna, 2016; Valentin-Vega et al., 2012). We previously published that increased uptake and consumption of glucose and glutamine, two carbon sources that are critically important for cancer cell metabolism, is in part due to inactivation of p53 and increased c-MYC stability downstream of ATM knockdown (Aird et al., 2015) and may be due to the regulation of glucose and glutamine transporters by these transcription factors (Schwartzenberg-Bar-Yoseph et al., 2004; Zhao et al., 2019). However, the transporter-dependence of nutrient uptake in ATM inhibited cells has never been directly tested. To answer this question, we used 3 ovarian cancer cell lines with functional/wildtype ATM and differential status of p53 and c-MYC (Ovcar8: mutant p53, c-MYC amplification; Ovcar3: mutant p53; Ovcar10: het mutant p53) and tested glucose and glutamine uptake upon inhibition of ATM. We found that pharmacological inhibition of ATM kinase activity using the small molecule inhibitor KU60019 increased glucose and glutamine consumption in all three cell lines (**Fig. 1A and Fig. S1A-C**). Consistently, a second ATM inhibitor AZD0156 also increased glucose and glutamine consumption (**Fig. 1B**). Suppression of ATM kinase activity was confirmed by decreased Chk2 phosphorylation (pChk2) (**Fig. 1C and Fig. S1D**). Interestingly, knockdown of the glucose transporters SLC2A1/GLUT1 and SLC2A4/GLUT4 or the glutamine transporter SLC1A5/ACST2 did not respectively alter the glucose or glutamine uptake induced by ATM inhibition (**Fig. 1D-G**), suggesting another route of metabolite uptake in these cells. Active ATM is known to signal through AKT to promote GLUT4 translocation (Halaby et al., 2008). Since we are inhibiting ATM, GLUT4 is not likely induced or translocated in our model. Together, these data suggest that inhibition of ATM increases uptake and consumption of multiple nutrients in a transporter-independent manner.

**Figure 1.**
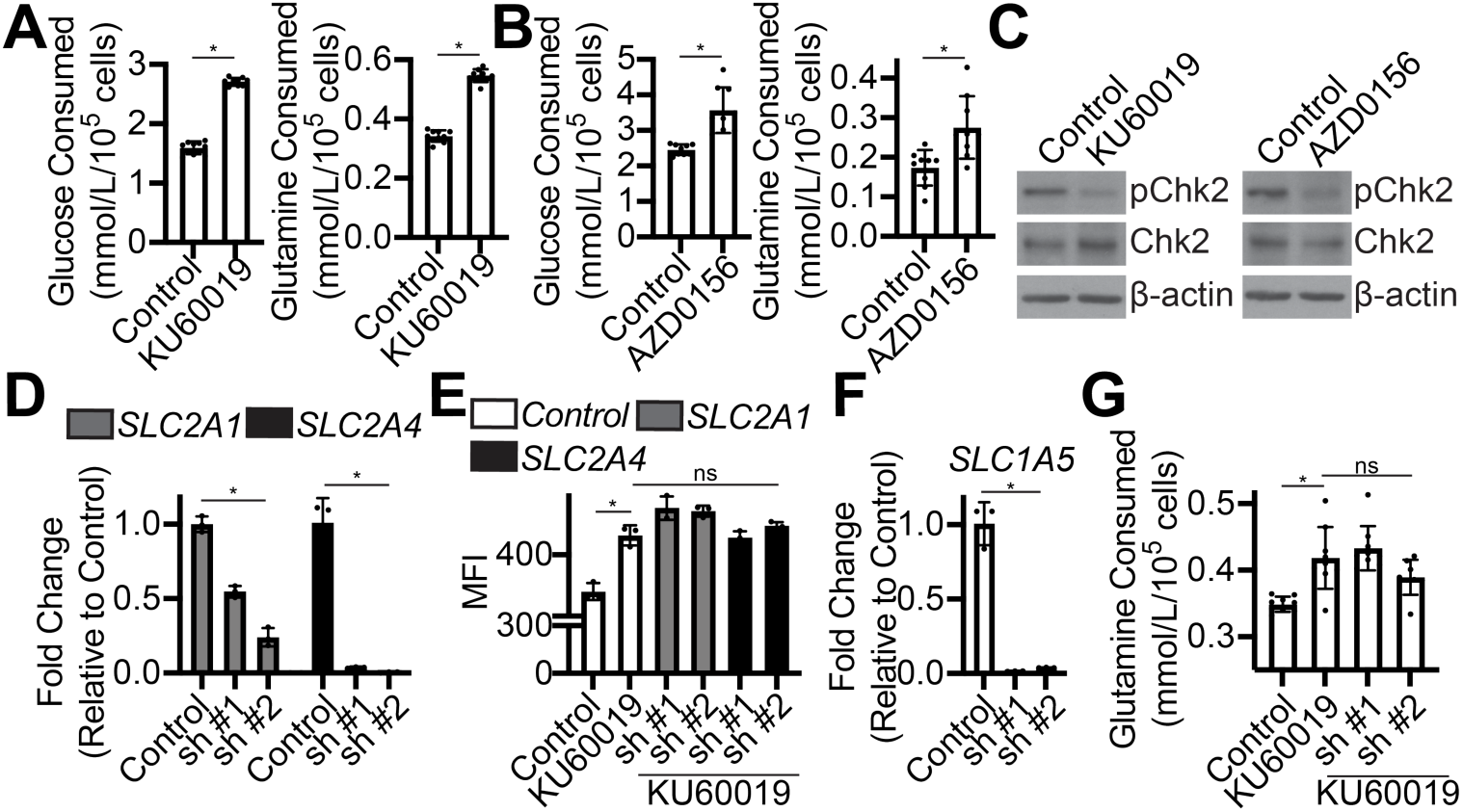
Knockdown of glucose and glutamine transporters in ATM inhibited cells does not affect glucose or glutamine consumption. **(A-C)** Ovcar8 cells were treated with the ATM inhibitors (KU60019 or AZD0156) for 24h. **(A-B)** Glucose and glutamine uptake were determined. n=9/group, one of 3 experiments is shown. Data represent mean ± SD. *p<0.005. **(C)** pChk2 and total Chk2 expression were determined by immunoblotting. β-actin was used as a loading control. One of 3 experiments is shown. **(D)** Ovcar3 cells were infected with lentivirus expressing short hairpin RNAs (shRNAs) targeting *SLC2A1* (GLUT1) or *SLC2A4* (GLUT4). shGFP was used as a control. mRNA expression was determined by RT-qPCR. n=3/group, one of 2 experiments is shown. Data represent mean ± SD. *p<0.01 **(E)** Same as (D), but cells were treated with KU60019 for 24h, and glucose uptake using the fluorescent glucose analog 2NBDG was determined by flow cytometry. MFI=median fluorescence intensity. n=3/group, one of 3 experiments is shown. Data represent mean ± SD. *p<0.0001; ns=not significant **(F)** Ovcar8 cells were infected with lentivirus expressing short hairpin RNAs (shRNAs) targeting *SLC1A5*. shGFP was used as a control. mRNA expression was determined by RT-qPCR. n=3/group, one of 2 experiments is shown. Data represent mean ± SD. *p<0.0001 **(G)** Same as (F), but cells were treated with KU60019 for 24h, and glutamine consumption was determined. n=3/group, one of 3 experiments is shown. Data represent mean ± SD. *p<0.005; ns=not significant

### Inhibition of ATM kinase activity induces macropinocytosis through activation of AMPK

Previous studies have shown that macropinocytosis is one mechanism whereby cancer cells scavenge nutrients from the microenvironment to support survival and proliferation (Commisso et al., 2013; Davidson et al., 2017; Hodakoski et al., 2019; Kamphorst et al., 2015; Kim et al., 2018; Lee et al., 2019; Palm, 2019; Palm et al., 2017; Recouvreux and Commisso, 2017; Redelman-Sidi et al., 2018; Yao et al., 2019; Zhang and Commisso, 2019). Due to its large size, fluorescently-labeled dextran is used as a surrogate for macropinocytosis as it cannot be taken up through other endocytic pathways (Commisso et al., 2014; Ivanov, 2008). Excitingly, suppression of ATM using multiple inhibitors or shRNA-mediated knockdown increased uptake of fluorescently-labeled dextran in multiple cell lines, indicating increased macropinocytosis in these cells (**Fig. 2A-D and Fig. S2A-C)**. Inhibition of ATM kinase activity was confirmed by decreased pChk2 (**Fig. S2D**). Ataxia telangiectasia and Rad3-related protein (ATR) is another kinase that is associated with the DNA damage response and has similar and often overlapping roles with ATM (Shiloh, 2003). Inhibition or knockdown of ATR, which decreases pChk1, did not increase dextran uptake, suggesting this is specific for ATM (**Fig. S2E-H**). This is consistent with our previous report demonstrating that knockdown of ATM, but not ATR, rescues proliferation defects due to metabolic deficiencies (Aird et al., 2015). To further confirm the increase in dextran uptake is due to macropinocytosis, we treated cells with EIPA, a Na+/H+ exchanger that is the gold standard inhibitor of macropinocytosis (Commisso et al., 2014; Ivanov, 2008). EIPA suppressed the increased dextran uptake in cells treated with the ATM inhibitor (**Fig. 2B-E**). EIPA in combination with the ATM inhibitor did not increase pChk2 (**Fig. S2I**), suggesting that increased ATM activity is not the cause of decreased macropinocytosis. EIPA alone decreased pChk2, although to varying degrees in the cell lines tested. This indicates that the signaling downstream of ATM to regulate macropinocytosis is likely not through Chk2, and pChk2 is only a surrogate for confirming activity of the ATM inhibitors. To validate our findings in a different model, we used fibroblasts from patients with ATM mutations who have a disorder termed ataxia-telangiectasia (A-T), who are predisposed to cancer and also exhibit metabolic dysfunction (McKinnon, 2012). A-T patient fibroblasts with various ATM mutations displayed increased dextran uptake compared to ATM wildtype IMR90 fibroblasts (**Fig. 2F**). Treatment of A-T patient fibroblasts or ATM inhibited IMR90 cells with EIPA decreased dextran uptake (**Fig. 2G and S2J-K**). Together, these data indicate that suppression of ATM kinase activity induces macropinocytosis.

**Figure 2.**
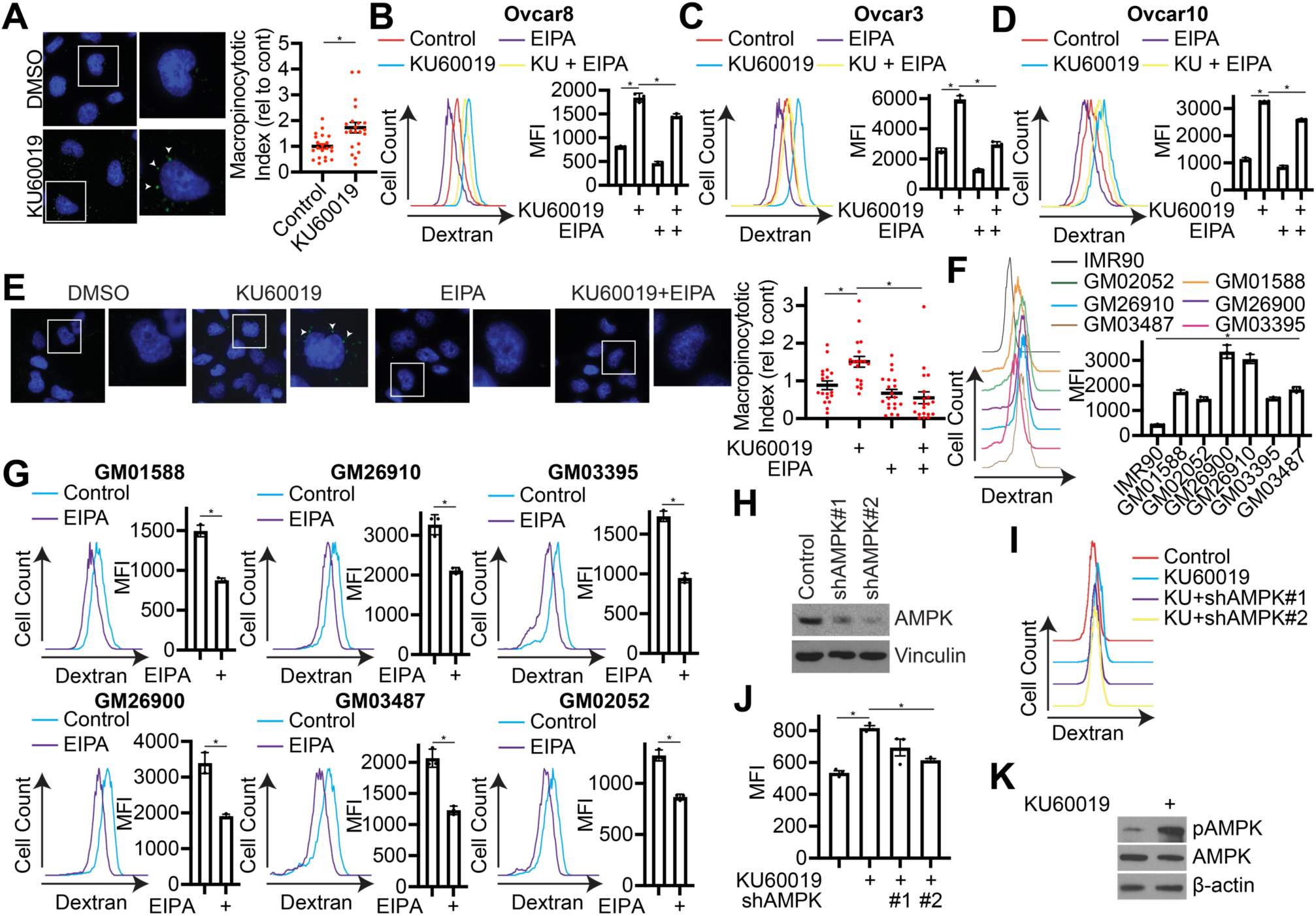
Inhibition of ATM kinase activity induces macropinocytosis through activation of AMPK. **(A)** Ovcar8 cells were treated with the ATM inhibitor KU60019 for 2h, and dextran uptake was determined by immunofluorescence. Arrowheads point to dextran-FITC. N>20/group, one of 3 experiments is shown. Data represent mean ± SD. *p<0.005 **(B-E)** Ovcar8 **(B, E)**, Ovcar3 **(C)**, and Ovcar10 **(D)** cells were treated with the ATM inhibitor KU60019 alone or in combination with the macropinocytosis inhibitor EIPA for 2h, and dextran uptake was determined by flow cytometry. MFI=median fluorescence intensity. n=3/group, one of at least 3 experiments is shown. Data represent mean ± SD. *p<0.0001 **(E)** Dextran uptake was determined by immunofluorescence. Arrowheads point to dextran-FITC. N>18/group, one of 2 experiments is shown. Data represent mean ± SD. *p<0.005 **(F)** Dextran uptake in IMR90 (ATM WT) and A-T fibroblasts was determined by flow cytometry. MFI=median fluorescence intensity. n=3/group, one of 3 experiments is shown. Data represent mean ± SD. *p<0.0001 **(G)** Dextran uptake in A-T patient fibroblasts treated with the macropinocytosis inhibitor EIPA for 2h was determined by flow cytometry. MFI=median fluorescence intensity. n=3/group, one of 3 experiments is shown. Data represent mean ± SD. *p<0.005 **(H-J)** Ovcar8 cells were infected with lentivirus expressing short hairpin RNAs (shRNAs) targeting AMPK. shGFP was used as a control. **(H)** Immunoblot analysis of AMPK. Vinculin was used as a loading control. One of 2 experiments is shown. **(I)** Dextran uptake was determined by flow cytometry. **(J)** Quantification of (I). MFI=median fluorescence intensity. n=3/group, one of 2 experiments is shown. Data represent mean ± SD. *p<0.05 **(K)** Ovcar8 cells were treated with the ATM inhibitor KU60019 for 2h. Immunoblot analysis of pAMPK and total AMPK. β-actin was used as a loading control. One of 5 experiments is shown.

Next we aimed to determine the mechanism of macropinocytosis induction downstream of ATM inhibition. Towards this goal, we used pharmacological or genetic inhibition of PI3K/AKT, WNT, and AMPK, three known inducers of macropinocytosis (Commisso et al., 2013; Kim et al., 2018; Redelman-Sidi et al., 2018; Tejeda-Munoz et al.). While inhibition of PI3K, AKT, or WNT had no effect (**Fig. S2L-M**), suppression of AMPK using shRNA decreased dextran uptake in ATM inhibited cells in a dose-dependent fashion (**Fig. 2H-J**). Consistently, we observed an increase in phosphorylation of AMPK upon inhibition of ATM in multiple cell lines (**Fig. 2K and S2N**). AMPK is phosphorylated by multiple kinases in response to nutrient stress and other stimuli (Herzig and Shaw, 2018). Previous reports have demonstrated that AMPK is both directly phosphorylated by ATM or its phosphorylation by LKB1 occurs in an ATM-dependent manner (Alexander et al., 2010; Luo et al., 2013; Sanli et al., 2010; Sanli et al., 2014; Sun et al., 2007; Suzuki et al., 2004; Tripathi et al., 2013), suggesting that direct or LKB1-mediated activation of AMPK is not occurring in ATM inhibited cells. Loss of ATM induces mitochondrial reactive oxygen species (ROS) (Valentin-Vega et al., 2012), and AMPK may be activated by mitochondrial ROS (Rabinovitch et al., 2017). This suggests that inhibition of ATM may increase macropinocytosis via mitochondrial ROS-mediated AMPK phosphorylation, although further experiments are needed to directly test this mechanism. Finally, we aimed to determine whether ATM inhibition-mediated macropinocytosis occurs through actin remodeling. AMPK can affect actin remodeling and macropinocytosis through the Pak1 pathway (Kim et al., 2018). We found that knockdown of Pak1 decreased macropinocytosis due to ATM inhibition (**Fig. S2O-P**), indicating this is the actin remodeling pathway that is likely induced downstream of ATM inhibition and AMPK phosphorylation. Together, these data suggest that AMPK activates macropinocytosis downstream of ATM inhibition in a Pak1-dependent manner.

### Suppression of macropinocytosis limits survival only in ATM inhibited cells

We next aimed to determine whether macropinocytosis is required for survival of ATM inhibited cells. We treated cells with the ATM inhibitor KU60019 and inhibited macropinocytosis using EIPA. The combination significantly decreased proliferation and increased apoptosis compared to single treatment controls in multiple cell lines (**Fig. 3A-B and S3A-B**). Similar results were observed *in vivo* where EIPA only affected ovarian tumor growth in combination with the ATM inhibitor KU60019 (**Fig. 3C-E**). *In vivo* the combination also induced apoptosis as we observed increased cleaved caspase 3 staining by IHC in the tumors (**Fig. 3F-G**). We also validated that KU60019 inhibited ATM by blotting for pChk2 in the tumors (**Fig. 3H and Fig. S3C**). Similar to our *in vitro* studies, we noted that pChk2 was decreased in the EIPA alone treated mice. To confirm inhibition of ATM induces macropinocytosis *in vivo*, mice were injected with fluorescently-labeled dextran 30min prior to harvesting the tumor. Consistent with our *in vitro* data (**Fig. 2**), KU60019 induced macropinocytosis, which was suppressed by EIPA (**Fig. 3I-J**). We observed co-localization of dextran with the lysosomal marker LAMP2, suggesting that the nutrients taken up by macropinocytosis are degraded within the lysosome. Together, these data demonstrate that inhibition of macropinocytosis limits proliferation/survival only of cells where ATM is inhibited and identify a metabolic vulnerability of tumors with low or mutated ATM.

**Figure 3.**
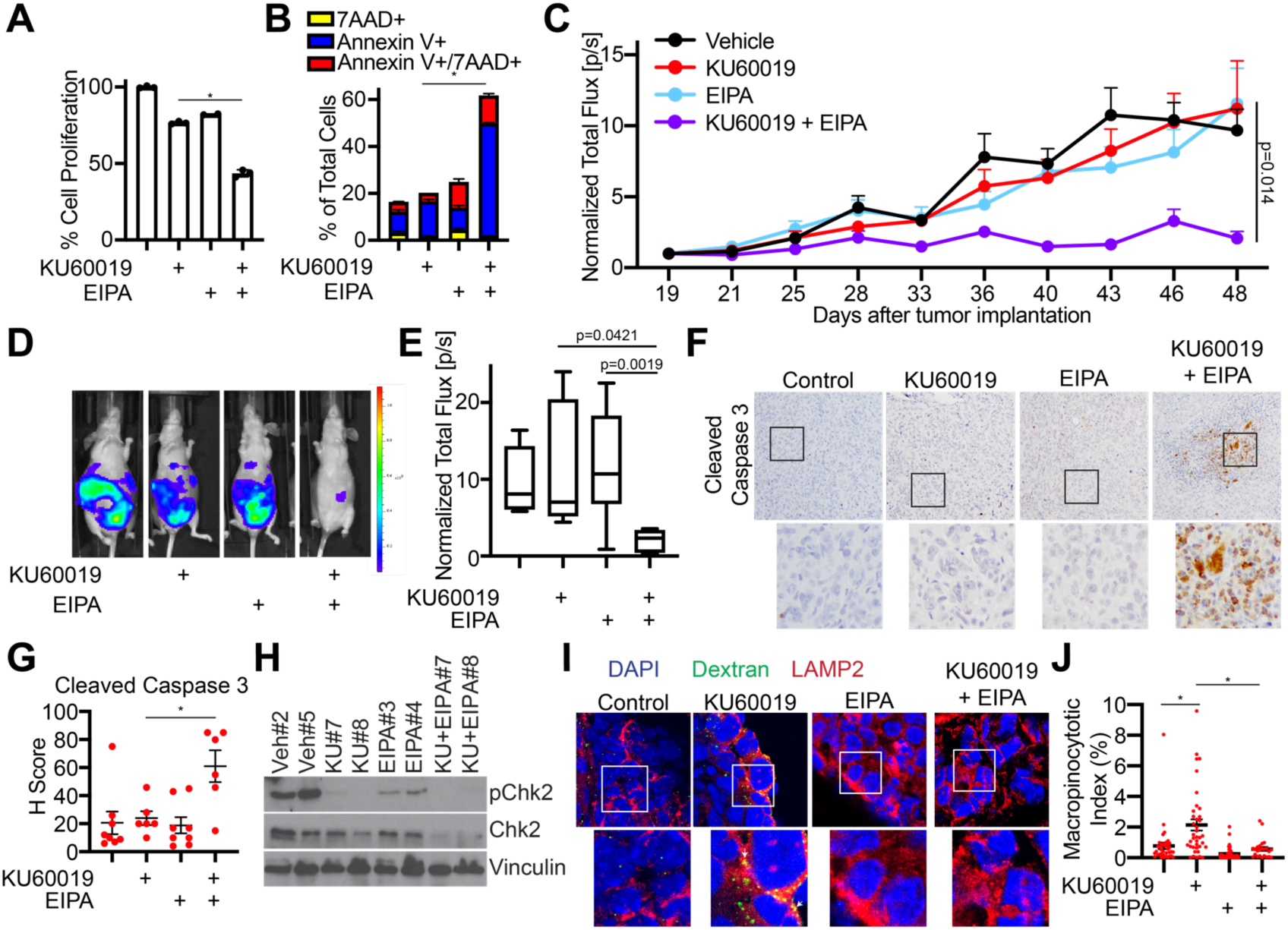
Inhibition of macropinocytosis is a vulnerability of ATM inhibited cells both *in vitro* and *in vivo*. **(A-B)** Ovcar8 cells were treated with the ATM inhibitor KU60019 or the macropinocytosis inhibitor EIPA alone or in combination for 4 days. **(A)** Proliferation was assessed by colony formation. n=3/group, one of 3 experiments is shown. Data represent mean ± SD. *p<0.001. **(B)** Apoptosis was assessed by Annexin V/7AAD staining. n=3/group, one of 3 experiments is shown. Data represent mean ± SD. *p<0.001 **(C)** Ovcar8 cells expressing luciferase were injected intraperitoneally into nude mice (6-8 mice/group). At day 19, mice were randomized and thereafter treated daily with the ATM inhibitor KU60019 or the macropinocytosis inhibitor EIPA alone and in combination. Shown is the tumor growth curve over 30 days normalized to Day 19 tumors. **(D)** Representative luciferase images of mice from each group at Day 48. **(E)** Quantification of (D) normalized to Day 19 tumors. **(F)** Representative cleaved caspase 3 immunohistochemistry in the indicated tumors. **(G)** H-Score of cleaved caspase 3 IHC. Data represent mean ± SEM. *p<0.01 **(H)** Representative pChk2 and total Chk2 immunoblot analysis of the indicated tumors. Vinculin was used as a loading control. **(I)** Representative confocal images of dextran-FITC and the lysosomal marker LAMP2 in the indicated tumors. Arrows point to dextran-FITC puncta co-localized with LAMP2. **(J)** Macropinocytotic index of dextran-FITC uptake in the tumors was assessed. Data represent mean ± SEM. *p<0.0005

### Macropinocytosis induced by ATM inhibition increases BCAA uptake to promote proliferation and survival

Finally, we aimed to determine the metabolic consequences of increased macropinocytosis due to inhibition of ATM. Previous studies have demonstrated that diverse nutrients including proteins, lipids, glucose, and others are taken up by macropinocytosis to promote cancer cell survival under nutrient stress (Commisso et al., 2013; Hodakoski et al., 2019; Kamphorst et al., 2015; Kim et al., 2018; Lee et al.; Palm, 2019; Palm et al., 2017; Recouvreux and Commisso, 2017; Redelman-Sidi et al., 2018; Tejeda-Munoz et al.; Zhang and Commisso, 2019). To determine the metabolites that may be important for tumors in the ATM inhibitor treated mice, we performed metabolomics on the tumor tissue, interstitial fluid, and the ascites fluid, which commonly occurs in ovarian cancer models. Interstitial fluid and ascites fluid can be used to assess microenvironment metabolite abundance. We observed an overall increase in multiple nucleotides within the tumors treated with the ATM inhibitor (**Fig. 4A and Table S1**), which is consistent with our previous report (Aird et al., 2015). We also observed an increase in metabolites related to the TCA cycle. In the combination group, TCA cycle metabolites and nucleotides were decreased, although nucleotides to a much lesser extent (**Fig. 4B**). This suggests that while ATM inhibition increases nucleotide metabolism, the TCA cycle and metabolites feeding into the TCA cycle are likely more important for survival and proliferation of ATM inhibited tumors. To investigate metabolite uptake from the microenvironment, we assessed metabolite abundance in the ascites and interstitial fluids. Valine and (iso)leucine, both branched chain amino acids (BCAAs), were decreased upon treatment with the ATM inhibitor and increased in the combination group (**Fig. 4C and Table S1**), suggesting that these carbon sources are consumed by ATM inhibited tumors to maintain proliferation and survival. Interestingly, BCAAs were not increased in the tumors, potentially because they have been catabolized for downstream pathways, such as the TCA cycle, or for protein synthesis. Finally, we assessed AMPK activation in the tumors. pAMPK was increased slightly in the ATM inhibitor only-treated group and further increased in the combination treated tumors (**Fig. S4A**). This suggests that the combination treatment is effective at least in part due to depletion of nutrients, leading to sustained AMPK activation.

**Figure 4.**
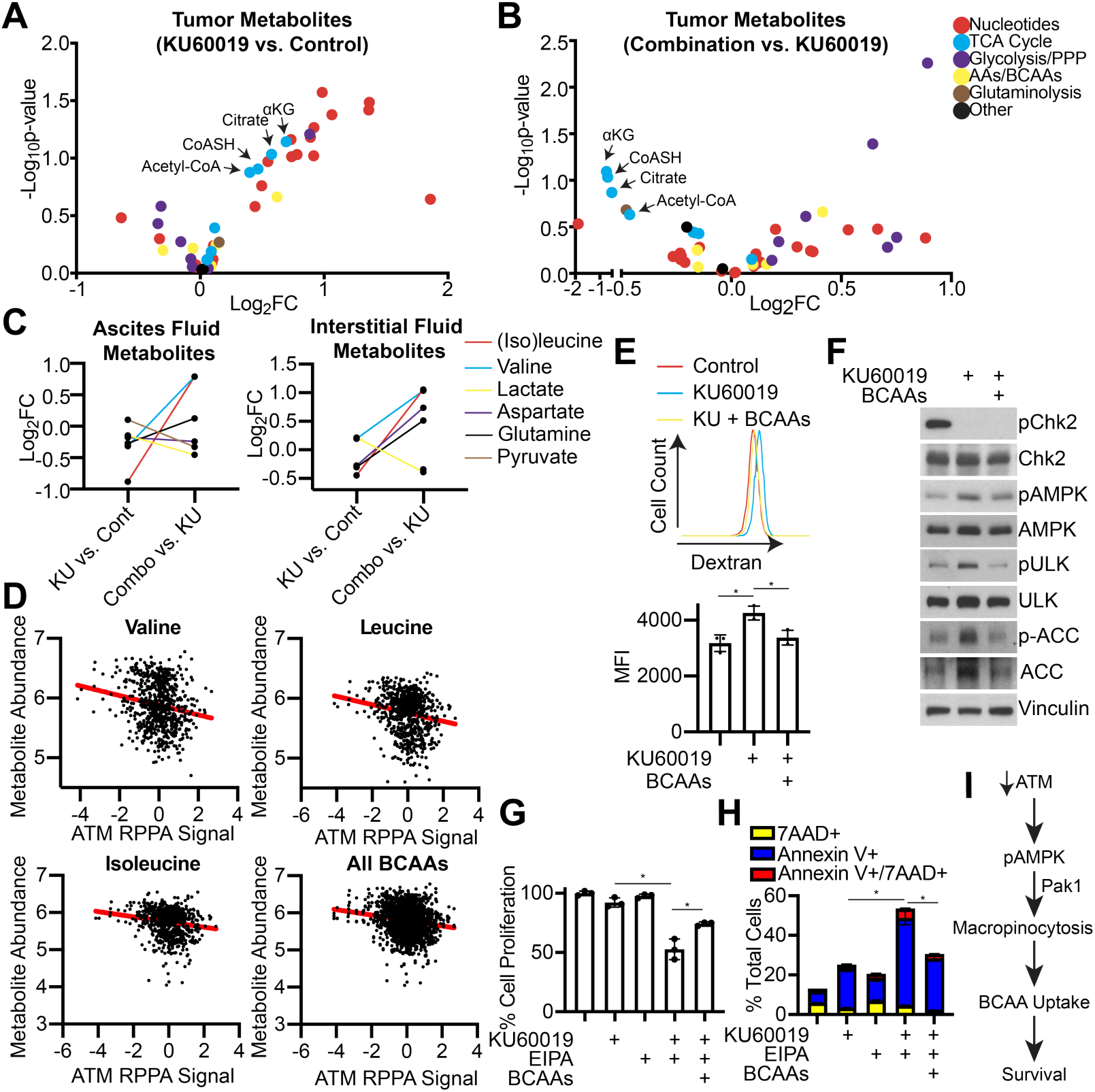
ATM inhibitor-induced macropinocytosis leads to increased branched chain amino acid uptake to promote cell survival. **(A)** Volcano plot of fold change (Log_2_FC) of metabolites in ATM inhibitor KU60019-treated vs. control tumors. **(B** Volcano plot of fold change (Log_2_FC) of metabolites in combination-treated (KU60019+EIPA) vs. KU60019-treated tumors. **(C)** Fold change (Log_2_FC) of metabolites in the ascites fluid and interstitial fluid in the indicated treatment groups. **(D)** BCAA metabolite abundance vs. ATM protein expression from DepMap.org. **(E)** Ovcar8 cells were treated for 2h with the ATM inhibitor KU60019 alone or in combination with BCAAs (mixture of valine, isoleucine, leucine). Dextran uptake was determined by flow cytometry. MFI=median fluorescence intensity. n=3/group, one of 3 experiments is shown. Data represent mean ± SD. *p<0.001 **(F)** Ovcar8 cells were treated for 2h with the ATM inhibitor KU60019 and a combination of BCAAs (mixture of valine, isoleucine, leucine), and expression of the indicated proteins was determined. Vinculin was used as a loading control. **(G)** Ovcar8 cells were treated with the ATM inhibitor KU60019 and a combination of EIPA or BCAAs (mixture of valine, isoleucine, leucine) for 4 days. Cell proliferation was assessed by colony formation. n=3/group, one of 3 experiments is shown. Data represent mean ± SD. *p<0.05 **(H)** Same as (F), but apoptosis was assessed by Annexin V/7AAD staining. n=3/group, one of 3 experiments is shown. Data represent mean ± SD. *p<0.0005 **(I)** Schematic of proposed model. Inhibition of ATM promotes macropinocytosis in an AMPK and Pak1-dependent manner to increase BCAAs and promote survival.

Next, we aimed to determine whether uptake of BCAAs in ATM inhibitor treated cells are critical for macropinocytosis and cancer cell survival *in vitro*. Interestingly, using publicly-available datasets from cell lines (depmap.org), we found protein expression of ATM or pChk2 negatively correlated with abundance of BCAAs and the TCA cycle metabolite citrate identified in our *in vivo* studies (**Fig. 4D and S4B-C**), suggesting this is a universal phenomenon of ATM low or inhibited cells. Notably, these studies were not done under nutrient-limiting conditions (Li et al., 2019), suggesting that there is basal macropinocytosis, which is likely further amplified during nutrient stress. If macropinocytosis increases BCAA uptake, we reasoned that supplementing cells with exogenous BCAAs would abrogate macropinocytosis in ATM inhibited cells. Indeed, supplementation of ATM inhibited cells with exogenous BCAAs suppressed macropinocytosis (**Fig. 4E**). Supplementation with BCAAs also abrogated the increase in AMPK phosphorylation and downstream signaling through ACC and ULK (**Fig. 4F**), suggesting that BCAA depletion is in part the cause of activated AMPK signaling. Consistent with the idea that BCAA uptake in ATM inhibited cells promotes proliferation and survival, supplementation of cells with BCAAs partially rescued the inhibition of proliferation and apoptosis observed in ATM inhibitor plus EIPA-treated cells (**Fig. 4G-H**). Together, these data suggest that inhibition of ATM increases macropinocytosis to promote cell survival at least in part through uptake and consumption of BCAAs from the microenvironment. The downstream mechanism of how BCAAs are affecting cell survival in ATM inhibited cells remains unknown. The irreversible step of BCAA metabolism via the branched chain alpha-keto acid dehydrogenase (BCKDH) complex generates mitochondrial NADH in addition to providing acetyl- and succinyl-CoA to the TCA cycle, and BCAAs have been shown to inhibit ROS (Sivanand and Vander Heiden, 2020). Since ATM inhibition increases ROS (Valentin-Vega et al., 2012), which in turn can promote AMPK activation (Rabinovitch et al., 2017), it is possible that BCAAs are consumed to promote redox homeostasis. Future studies are warranted to dissect this mechanism.

In summary, we identified a new mechanism of macropinocytosis through inhibition of ATM, which increases uptake of BCAAs to promote cancer cell proliferation and survival under nutrient stress (**Fig. 4I**). This study provides a novel mechanism of metabolism controlled by ATM inhibition and suggests that macropinocytosis or BCAA metabolism represents a novel metabolic vulnerability that can be targeted in combination with ATM inhibitors, which have recently entered clinical trials (clinicaltrials.gov). Moreover, this pathway may also be a vulnerability of ATM mutated or low expressing tumors. Since A-T patients with ATM mutations also have metabolic disorders, our data may have implications beyond cancer for controlling metabolism in the context of low or mutated ATM.

## Supplemental Figures

**Figure S1.**
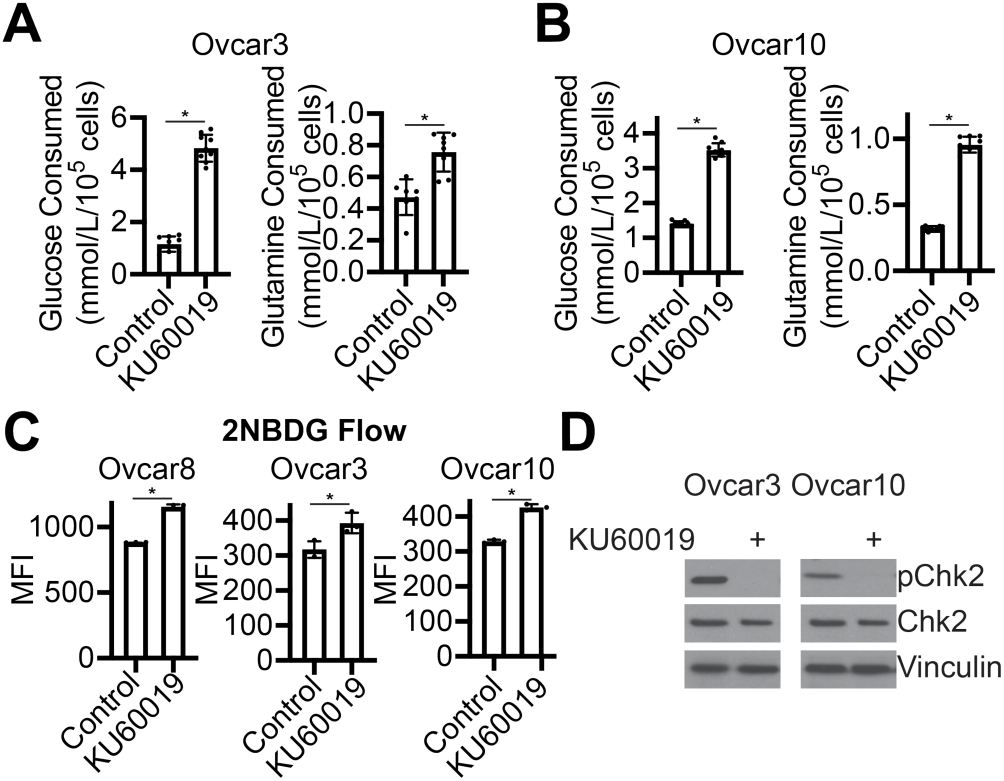
Inhibition of ATM increases glucose and glutamine consumption in multiple cell lines; related to Figure 1. **(A-B)** Ovcar3 (A) and Ovcar10 (B) cells were treated with the ATM inhibitor KU60019 for 24h. Glucose and glutamine consumption was determined. n=9/group, one of 2 experiments is shown. Data represent mean ± SD. *p<0.005 **(C)** Glucose uptake in the indicated cells by 2NBDG was determined by flow cytometry after treatment with the ATM inhibitor KU60019 for 24h. MFI=median fluorescence intensity. n=3/group, one of at least 2 experiments is shown. Data represent mean ± SD. *p<0.05 **(D)** The indicated cells were treated with KU60019 for 24h, and pChk2 and total Chk2 expression were determined by immunoblotting. Vinculin was used as a loading control. One of 3 experiments is shown

**Figure S2.**
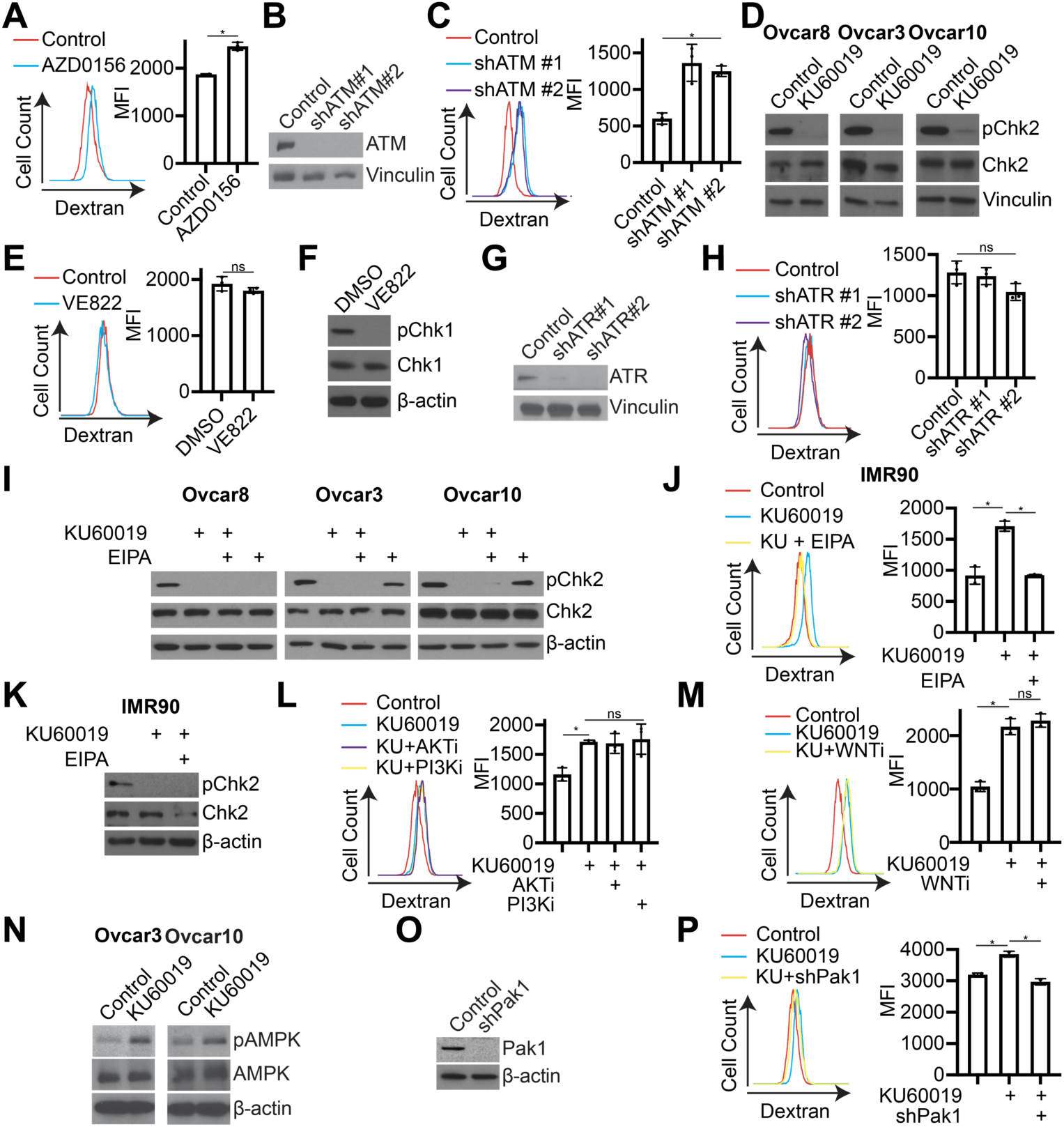
Inhibition of ATM, but not ATR, induces macropinocytosis and is independent of PI3K and WNT; related to Figure 2. **(A)** Ovcar8 cells were treated with the ATM inhibitor AZD0156 for 2h, and dextran uptake was determined by flow cytometry. MFI=median fluorescence intensity. n=3/group, one of 3 experiments is shown. Data represent mean ± SD. *p<0.005 **(B-C)** Ovcar8 cells were infected with lentivirus expressing short hairpin RNAs (shRNAs) targeting ATM. shGFP was used as a control. **(B)** Immunoblot analysis of ATM. Vinculin was used as a loading control. One of 3 experiments is shown. **(C)** Dextran uptake was determined by flow cytometry. MFI=median fluorescence intensity. n=3/group, one of 3 experiments is shown. Data represent mean ± SD. *p<0.0001 vs. control **(D)** Ovcar8, Ovcar3, and Ovcar10 cells were treated with the ATM inhibitor KU60019 for 2h. Immunoblot analysis of p-Chk2 and total Chk2. Vinculin was used as a loading control. One of 3 experiments is shown. **(E-F)** Ovcar8 cells were treated with the ATR inhibitor VE822. **(E)** Dextran uptake was determined by flow cytometry. MFI=median fluorescence intensity. n=3/group, one of 3 experiments is shown. Data represent mean ± SD. ns=not significant. **(F)** Immunoblot analysis of p-Chk1 and total Chk1. β-actin was used as a loading control. One of 3 experiments is shown. **(G-H)** Ovcar8 cells were infected with lentivirus expressing short hairpin RNAs (shRNAs) targeting ATR. shGFP was used as a control. **(G)** Immunoblot analysis of ATR. Vinculin was used as a loading control. One of 3 experiments is shown. **(H)** Dextran uptake was determined by flow cytometry. MFI=median fluorescence intensity. n=3/group, one of 3 experiments is shown. Data represent mean ± SD. ns=not significant. **(I)** Ovcar8, Ovcar3, and Ovcar10 cells were treated with the ATM inhibitor KU60019 alone or in combination with EIPA for 2h. Immunoblot analysis of pChk2 and total Chk2. β-actin was used as a loading control. One of 3 experiments is shown. **(J-K)** IMR90 fibroblasts (ATM WT) were treated with the ATM inhibitor KU60019 or in combination with EIPA for 2h. **(J)** Dextran uptake was determined by flow cytometry. MFI=median fluorescence intensity. n=3/group, one of 3 experiments is shown. Data represent mean ± SD. *p<0.0001. **(K)** Immunoblot analysis of pChk2 and total Chk2. β-actin was used as a loading control. One of 3 experiments is shown. **(L)** Ovcar8 cells were treated with the ATM inhibitor KU60019 alone or in combination with the AKT inhibitor AZD5363 or the PI3K inhibitor buparisib for 2h. Dextran uptake was determined by flow cytometry. MFI=median fluorescence intensity. n=3/group, one of 2 experiments is shown. Data represent mean ± SD. *p<0.05 **(M)** Ovcar8 cells were treated with the ATM inhibitor KU60019 alone or in combination with the WNT inhibitor ICG-001 for 2h. Dextran uptake was determined by flow cytometry. MFI = median fluorescence intensity. n=3/group, one of 2 experiments is shown. Data represent mean ± SD. *p<0.0001 ns=not significant **(N)** Ovcar3 or Ovcar10 cells were treated with the ATM inhibitor KU60019 for 2h. Immunoblot analysis of pAMPK and AMPK. β-actin was used as a loading control. One of 3 experiments is shown. **(O-P)** Ovcar8 cells were infected with lentivirus expressing a short hairpin RNA (shRNA) Pak1. shGFP was used as a control. **(O)** Immunoblot analysis of Pak1. β-actin was used as a loading control. One of 2 experiments is shown. **(P)** Dextran uptake was determined by flow cytometry. MFI=median fluorescence intensity. n=2-3/group, one of 2 experiments is shown. Data represent mean ± SD. *p<0.01

**Figure S3.**
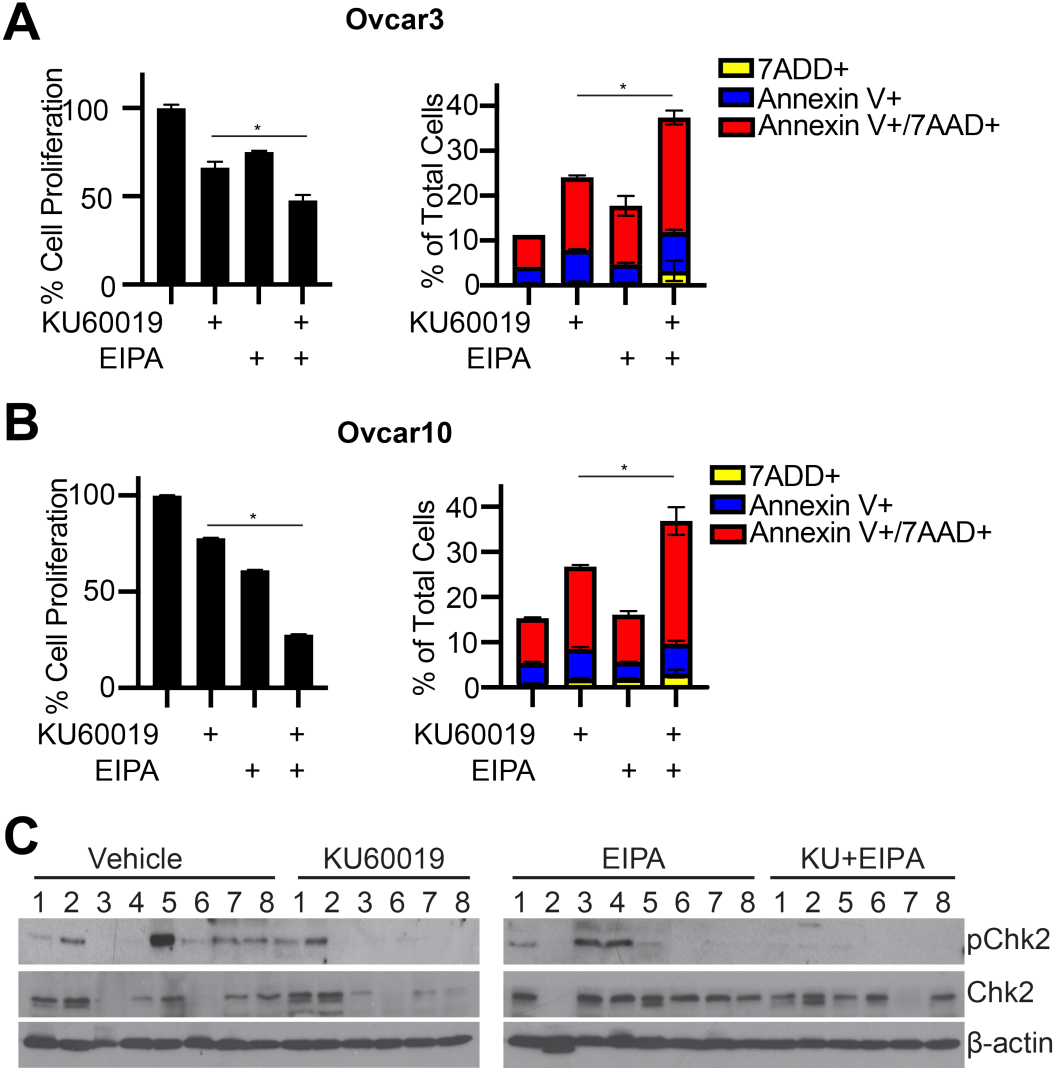
Inhibition of macropinocytosis inhibits cell proliferation and increases cell death in ATM inhibitor-treated cells. Related to Figure 3. **(A-B)** Ovcar3 (A) or Ovcar10 (B) cells were treated with the ATM inhibitor KU60019 or the macropinocytosis inhibitor EIPA alone and in combination for 4 days. Proliferation (left panel) was assessed by colony formation. Apoptosis (right panel) was assessed by Annexin V/7AAD staining. n=3/group, one of 3 experiments is shown. Data represent mean ± SD. *p<0.01 **(C)** pChk2 and total Chk2 immunoblot analysis in all of the tumors. β-Actin was used as a loading control.

**Figure S4.**
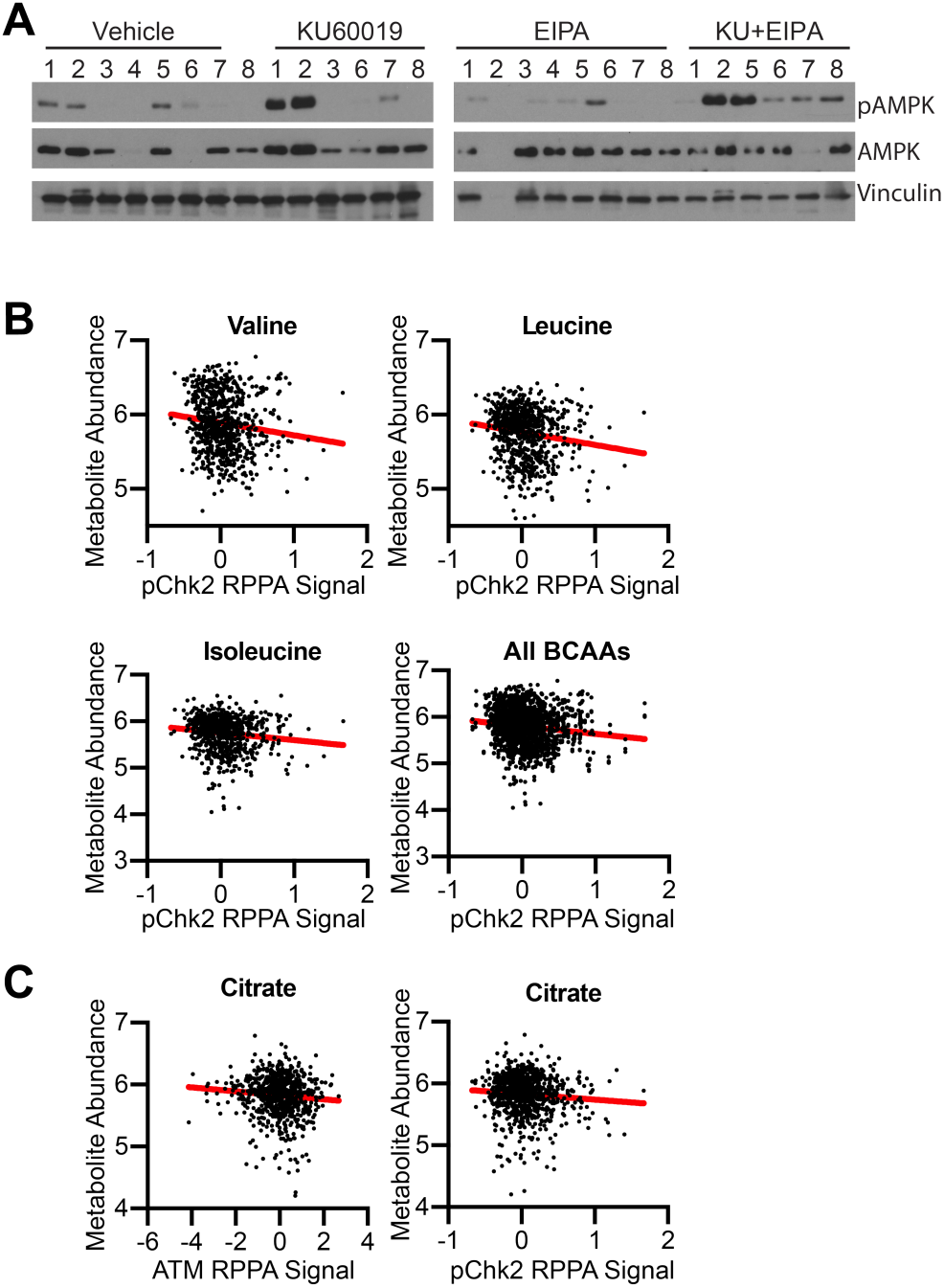
ATM and pChk2 protein expression negatively correlate with branched chain amino acids and the TCA cycle metabolite citrate. Related to Figure 4. **(A)** AMPK and total AMPK immunoblot analysis in all of the tumors. Vinculin was used as a loading control. **(B)** BCAAs metabolite abundance vs. pChk2 protein expression from DepMap.org. **(C)** Citrate metabolite abundance vs. ATM or pChk2 protein expression from DepMap.org.

## Acknowledgments

We would like to acknowledge Dr. Naveen Kumar Tangudu for providing helpful discussion. This work was supported by grants from the National Institutes of Health (F31CA236372 to E.S.D., F31CA250366 to K.E.L., R01GM132261 to N.W.S., and R37CA240625 and R00CA194309 to K.M.A.), the Congressionally Directed Medical Research Program (W81XWH-18-1-0103 to K.M.A.), and a Penn State Cancer Institute Postdoctoral Fellowship (R.B.).

## Author Contributions

Conceptualization, C.W.C. and K.M.A.; Methodology, E.L.V., E.V.K., N.W.S.; Investigation, C.W.C, R.B., E.S.D., K.E.L., E.L.V., E.V.K, N.W.S., K.M.A.; Writing, C.W.C., N.W.S, and K.M.A.; Visualization, C.W.C, N.S.W., K.M.A.; Supervision, N.W.S., K.M.A.; Funding Acquisition, N.W.S., and K.M.A.

## Materials and Methods

### Cell lines and culture conditions

Ovcar8, Ovcar3, and Ovcar10 cells were cultured in RPMI (Corning, Cat# 10-040-CV) with 5% FBS. 293FT cells were cultured in DMEM (Corning, Cat# 10-013-CV) with 5% FBS. Normal diploid IMR90 (purchased from ATCC) and A-T human fibroblasts (GM01588, GM02052, GM03995, GM04387, GM22690, and GM22691 purchased from Coriell Institute) were cultured in low oxygen (2% O_2_) in DMEM (Corning, Cat# 10-017-CV) with 15% FBS supplemented with L-glutamine, non-essential amino acids, sodium pyruvate, and sodium bicarbonate. Experiments were performed on IMR90 between population doubling #25-35 and A-T fibroblasts below passage 10. All cell lines were cultured in MycoZap and were routinely tested for mycoplasma using a highly sensitive PCR-based method (Uphoff and Drexler, 2005). Tumor cell lines were authenticated using STR Profiling using Genetica DNA Laboratories.

### Glucose uptake by flow cytometry

Cells (10^5^/well in 12-well plates) were incubated with 5µM of the fluorescent glucose analog 2NBDG (N13195, Thermo Fisher Scientific) in Opti-MEM™ I Reduced Serum Medium (31985088, Thermo Fisher Scientific) for 2h. Cells were run on a 10-color FACSCanto flow cytometer (BD Biosciences). Data were analyzed with FlowJo Software.

### YSI Bioanalyzer Analysis

Glucose and glutamine consumption were measured using a YSI 2950 Bioanalyzer (Yellow Springs, OH). Briefly, the same number of cells (10^5^/well in 12-well plates) were seeded in complete media. One day later, the media was changed to RPMI+0.1% FBS and cells were incubated in 10μM KU60019 or 1μM AZD0156 for 24h. Media was collected, and the number of cells per well was counted to normalize for cell number.

### Western blotting

Cell lysates were collected in 1X sample buffer (2% SDS, 10% glycerol, 0.01% bromophenol blue, 62.5mM Tris, pH 6.8, 0.1M DTT) and boiled (10 min at 95°C). Protein concentration was determined using the Bradford assay. Proteins were resolved using SDS-PAGE gels and transferred to nitrocellulose membranes (Fisher Scientific) (110mA for 2 h or overnight at 4°C). Membranes were blocked with 5% nonfat milk or 4% BSA in TBS containing 0.1% Tween-20 (TBS-T) for 1 h at room temperature. Membranes were incubated overnight at 4°C in primary antibodies (**Table S2**) in 4% BSA/TBS + 0.025% sodium azide. Membranes were washed 4 times in TBS-T for 5 min at room temperature after which they were incubated with HRP-conjugated secondary antibodies (Cell Signaling, Danvers, MA) for 1 h at room temperature. After washing 4 times in TBS-T for 5 min at room temperature, proteins were visualized on film after incubation with SuperSignal West Pico PLUS Chemiluminescent Substrate (ThermoFisher, Waltham, MA).

### Lentiviral packaging and infection

Lentiviral constructs were transfected into 293FT cells using polyethylenimine (PEI). Lentivirus was packaged using the ViraPower Kit (Invitrogen, Carlsbad, CA, USA) following the manufacturer’s instructions. Cells were infected overnight with lentivirus targeting the gene of interest or control shGFP and selected with 1μg/ml puromycin for 3 days. The following shRNAs were used: shATM#1: TRCN0000038658; shATM#2: TRCN0000010299; shATR#1: TCRN0000039615; shATR#2: TCRN0000039616; shSLC2A1#1: TRCN0000043585; shSLC2A1#2: TRCN0000043584; shSLC2A4#1: TRCN0000043628; shSLC2A4#2; TRCN0000043629; shSLC1A5#1: TRCN0000043118; shSLC1A5#2: TRCN0000043119; shPAK1#1: TRCN0000002226; shAMPK#1: TRCN0000000857; shAMPK#2: TRCN0000000859.

### RT-qPCR

Total RNA was extracted from cells with Trizol, DNase treated, cleaned, and concentrated using Zymo columns (Zymo Research, Cat# R1013) following the manufacturer’s instructions. Optical density values of RNA were measured using NanoDrop One (Thermo Scientific) to confirm an A260 and A280 ratio above 1.9. Relative expression of target genes (**Table S3**) were analyzed using the QuantStudio 3 Real-Time PCR System (Thermo Fisher Scientific) with clear 96 well plates (Greiner Bio-One). Primers were designed using the Integrated DNA Technologies (IDT) tool (http://eu.idtdna.com/scitools/Applications/RealTimePCR/). A total of 25ng of RNA was used for One-Step qPCR (Quanta BioSciences) following the manufacturer’s instructions in a final volume of 10μl. Conditions for amplification were: 10 min at 48°C, 5 min at 95°C, 40 cycles of 10 s at 95°C and 7 s at the corresponding annealing temperature (**Table S3**). The assay ended with a melting curve program: 15 s at 95°C, 1 min at 70°C, then ramping to 95°C while continuously monitoring fluorescence. Each sample was assessed in triplicate. Relative quantification was determined to multiple reference genes (*B2M, MRPL9, PSMC4*, and *PUM1*) using the delta-delta Ct method.

### Flow cytometry

For Dextran-FITC experiments: cells were incubated with 1mg/mL dextran-FITC (Sigma-Aldrich, Cat # 90718-1G) in base media (Sigma-Aldrich, Cat# D5030) for 2h. Where indicated, cells were incubated with 10μM KU60019, 25μM EIPA, 4mM BCAAs (mixture of valine, isoleucine, and leucine), 10μM AZD5363, 10μM Buparisib, or 25μM ICG001 for 2h. After washing twice with 1X PBS, cells were trypsinized and run on a 10-color FACSCanto flow cytometer (BD Biosciences). For Annexin V/7AAD experiments: Cells (2×10^4^/well in 12-well plates) were treated for 4 days with 2μM KU60019, 2μM EIPA, and BCAAs (mixture of 4mM valine, isoleucine, leucine) in RPMI + 5% FBS. Cells were stained with 2×10^6^ cells/ml Annexin V (R37176, Thermo Fisher Scientific) and 0.5μg/ml 7AAD (13-6993-T200, Tonbo Biosciences) in 2.5 mM Ca2+-containing RPMI for 15 minutes at room temperature. Data were analyzed using FlowJo software (Ashland, OR).

### Dextran uptake by immunofluorescence

Cells were incubated with 5mg/ml dextran-FITC (Sigma-Aldrich, Cat# 90718-1G) in base media (D5030, Sigma) for 2h. A low pH buffer (0.1 M Sodium acetate + 0.05 NaCl) was used to wash away non-specific binding of dextran to the cell membrane. Cells were fixed in 4% paraformaldehyde (10min) and permeabilized in 0.2% Triton X-100 (5 min). Cells were further incubated with 0.15 μg/ml DAPI in PBS (1 min), mounted, and sealed. For macropinocytotic index, images were acquired at room temperature using a Nikon Eclipse 90i microscope with a 20x/0.17 objective (Nikon DIC N2 Plan Apo) equipped with a CoolSNAP Photometrics camera. Macropinocytotic index was determined using the method described in (Lee et al., 2019). For the tumor model: 0.2mg/mice Dextran, Oregon Green™ (D7173, Thermo Fisher Scientific) in 200μl Evans Blue dye (1% in sterile water; E2129, Sigma) was injected intraperitoneally 30min prior to harvesting tumors. Tumors were frozen at −80°C in OCT (Tissue Tek) and sectioned using a cryostat. Sections were stained with LAMP2 (1:100, Santa Cruz, Cat# sc-18822) in 3% BSA/PBS at room temperature for 1 h. Tissues were washed three times and then incubated in Cy3 anti-mouse secondary antibody (1:5000, Jackson ImmunoResearch Labs, Cat# 715-165-150) in 3% BSA/PBS at room temperature for 1 h. Finally, tissues were incubated with 0.15 μg/ml DAPI in PBS for 1min, washed three times with PBS, mounted and sealed. Images were acquired at room temperature using a confocal microscope (Leica SP8) with a 64X oil objective. Macropinocytotic index was determined using the method described in (Commisso et al., 2014).

### Colony Formation

Cells (10^5^/well in 12-well plates) were treated with the 2µM KU60019 and a combination of 2µM EIPA or 4mM BCAAs (mixture of valine, isoleucine, and leucine) in 5% FBS RPMI for 4 days. KU60019, EIPA, and BCAAs were supplied daily. Colony formation was visualized by fixing cells in 1% paraformaldehyde (5 min) and staining with 0.05% crystal violet (20 min). Wells were destained for 5 min in 500 mL 10% acetic acid. Absorbance (590nm) was measured using a spectrophotometer (Spectra Max 190). Each sample was assessed in triplicate.

### Murine tumor model

Two-month old female nude mice were purchased from Jackson Labs. All mice were maintained in a HEPA-filtered ventilated rack system at the Milton S. Hershey Medical Center animal facility. Mice were housed up to 5 mice per cage and in a 12-hour light/dark cycle. All experiments with animals were performed in accordance with institutional guidelines approved by the Institutional Animal Care and Use Committee (IACUC) at the Penn State College of Medicine. Ovcar8 ovarian cancer cells (5×10^6^ in 200μl PBS) were injected intraperitoneally into mice. Mice were monitored every 3-4 days by non-invasive luciferase imaging by intraperitoneal injection of 150 mg/kg Luciferin (PerkinElmer) and quantification of luciferase activity using Imaging Systems (IVIS Spectrum System; Xenogen Corporation). On day 19, mice were randomized and thereafter treated daily with 10mg/kg KU60019, 10mg/kg EIPA both alone and in combination via intraperitoneal injection daily. All animals were euthanized at day 48 post tumor implantation, and ascites fluid, interstitial fluid (Sullivan et al., 2019), and tumor tissues were collected.

### Immunohistochemistry

Immunohistochemistry (IHC) was conducted on formalin fixed paraffin embedded tissues using cleaved caspase 3 antibody (1:300, Cell Signaling, Cat# 9661), EnVision+ Dual Link HRP secondary (Agilent, Cat#K406311-2), and the ImmPACT DAB Peroxidase (HRP) Substrate (Vector Laboratories, Cat# SK-4105) following the manufacturer’s instructions. Briefly, the tissue was rehydrated by incubating in a series of zylenes/ethanol baths. Antigen retrieval was performed by steaming in citrate buffer (Thermo Fisher Scientific, Cat# 005000) for 40min. Endogenous peroxidase was quenched by incubating in 3% H_2_O_2_/MeOH for 20min after which slides were washed and blocked in 1% BSA/PBS at room temperature for 30min. Slides were incubated overnight at 4°C in primary antibody (1:300), washed, and incubated in secondary-HRP for 45min. DAB was used to develop, and Mayer’s Hemotoxylin (Sigma-Aldrich, Cat# MHS16) was used to counterstain the tissue. Finally, slides were dehydrated and mounted. H score was determined on blinded samples, where H score = [1 × (% cells 1+) + 2 × (% cells 2+) + 3 × (% cells 3+)].

### Metabolomics

Metabolomic profiling was conducted using ion pairing reversed phase liquid chromatography-high resolution mass spectrometry (LC-HRMS) modified from previous methods for polar analytes (Guo et al., 2016) and nucleotides (Kuskovsky et al., 2019). Samples were spiked with a stable isotope mix containing AMP-^13^C_10_,^15^N_5_, dAMP-^13^C_10_,^15^N_5_, ATP-^13^C_10_,^15^N_5_, dATP-^13^C_10_,^15^N_5_, dTMP-^13^C_10_,^15^N_2_, dTTP-^13^C_10_,^15^N_2_, dCMP-^13^C_9_,^15^N_3_, CTP-^13^C_9_,^15^N_3_, dCTP-^13^C_9_,^15^N_3,_ ^13^C_3_-Sodium Pyruvate, ^13^C_3_-Lactate, ^13^C_4_-Fumaric Acid, ^13^C_4_-Succinic acid, ^13^C_4_ ^15^N_1_-Aspartic Acid, ^13^C_6_-Citric Acid, ^13^C_6_-Glucose-6-phosphate, ^13^C_2_-AcetylCoA, ^13^C_5_-D-α-Hydroxyglutaric acid, ^13^C_5_ ^15^N_2_-Glutamine from Sigma-Aldrich or Cambridge Isotope Laboratories. 1 mL of Optima LC-MS grade 80:20 methanol:water prechilled to −80°C was then added to each sample, followed by a 30 second vortex mixing, 15 second pulse sonication with a probe tip sonicator then samples were returned to the −80 °C freezer for 30 min. Insoluble debris was precipitated by centrifugation for 10 min, 17,000 x g at 4°C. Supernatant was evaporated to dryness under nitrogen, resuspended in 100 µL 5% 5-sulfosaliclyic acid in water, and 5 µL was injected for analysis. LC-HRMS was conducted on an Ultimate 3000 UHPLC equipped with a refrigerated autosampler (at 6 °C) and a column heater (at 55 °C) with a HSS C18 column (2.1 × 100 mm i.d., 3.5 μm; Waters, Milford, MA) used for separations. Solvent A was 5 mM DIPEA and 200 mM HFIP and solvent B was methanol with 5 mM DIPEA 200 mM HFIP. The gradient was as follows: 100 % A for 3 min at 0.18 mL/min, 100 % A at 6 min with 0.2 mL/min, 98 % A at 8 min with 0.2 mL/min, 86 % A at 12 min with 0.2 mL/min, 40 % A at 16 min and 1 % A at 17.9 min-18.5 min with 0.3 mL/min then increased to 0.4 mL/min until 20 min. Flow was ramped down to 0.18 mL/min back to 100 % A over a 5 min re-equilibration. For MS analysis, the UHPLC was coupled to a Q Exactive HF mass spectrometer (Thermo Scientific, San Jose, CA, USA) equipped with a HESI II source operating in negative mode. The operating conditions were as follows: spray voltage 4000 V; vaporizer temperature 200 °C; capillary temperature 350 °C; S-lens 60; in-source CID 1.0 eV, resolution 60,000. The sheath gas (nitrogen) and auxiliary gas (nitrogen) pressures were 45 and 10 (arbitrary units), respectively. Single ion monitoring (SIM) windows were acquired around the [M-H]^−^ of each analyte with a 20 *m/z* isolation window, 4 *m/z* isolation window offset, 1e^6^ ACG target and 80 ms IT, alternating in a Full MS scan from 70-950 *m/z* with 1e6 ACG, and 100 ms IT. Data was analyzed in XCalibur v4.0 and/or Tracefinder v4.1 (Thermo) using a 5 ppm window for integration of the peak area of all analytes and internal standards used for normalization.

### Quantification and Statistical Analysis

GraphPad Prism version 8.0 was used to perform statistical analysis. One-way ANOVA or t-test were used as appropriate to determine p values of raw data. P-values < 0.05 were considered significant. Longitudinal and cross-sectional analysis of tumor volume where calculated using TumorGrowth tool using default parameters (Enot et al., 2018).

